# Low cost and sustainable hyaluronic acid production in a manufacturing platform based on *Bacillus subtilis* 3NA strain

**DOI:** 10.1101/2021.01.18.427161

**Authors:** Sebastián Cerminati, Mélanie Leroux, Pablo Anselmi, Salvador Peirú, Juan C. Alonso, Bernard Priem, Hugo G. Menzella

## Abstract

Hyaluronic acid (HA) is a high value glycosaminoglycan mostly used in health and cosmetic applications. Commercial HA is produced from animal tissues or in toxigenic bacteria of the genus *Streptococcus* grown in complex media, which are expensive and raise environmental concerns due to the disposal of large amounts of broth with high organic loads. Other microorganisms were proposed as hosts for the heterologous production of HA, but the methods are still costly. The extraordinary capacity of this biopolymer to bind and retain water attracts interest for large scale applications where biodegradable materials are needed, but its high cost and safety concerns are barriers for its adoption.

*Bacillus subtilis* 3NA strain is prototrophic, amenable for genetic manipulation, GRAS, and can rapidly reach high cell densities in salt-based media. These phenotypic traits were exploited to create a platform for biomolecule production using HA as a proof of concept. First, the 3NA strain was engineered to produce HA; second, a chemically defined medium was formulated using commodity-priced inorganic salts combined at the stoichiometric ratios needed to build the necessary quantities of biomass and HA; and third, a scalable fermentation process, where HA can be produced at the maximum volumetric productivity (VP), was designed.

A comparative economic analysis against other methods indicates that the new process may increase the operating profit of a manufacturing plant by more than 100 %. The host, the culture medium, and the rationale employed to develop the fermentation process described here, introduce an IP free platform that could be adaptable for production of other biomolecules.

**Key Points:**

- A platform for the production of biomolecules was designed based on *B. subtilis* 3NA, a chemically defined medium and a fermentation process.
- As proof of concept, high quality hyaluronic acid was produced with an environmentally friendly process.
- A techno-economic analysis indicates that the process is more that 100% profitable than current methods.

## Introduction

Hyaluronic acid (HA), also termed hyalunoran, is a linear glycan chain consisting of successive units of a disaccharide formed by β-1,3-D-glucuronic acid (GlcUA) and β-1,4-N-acetyl-D-glucosamine (GlcNAc) (Figure 1A) with molecular weights of up to 8 MDa (Weissmann and Meyer 1954). This polysaccharide, which is ubiquitous in vertebrates and in human tissues, is found in the extracellular matrix playing structural and physiological functions. It is also present in few selected microorganisms, forming the capsule of some pathogenic bacteria. Due to the presence of abundant polar groups, which bind water to form a hydrated gel, HA is an excellent moisturizing factor. HA diverse properties, along with its lack of immunogenicity, made this polysaccharide attractive for cosmetic and pharmaceutical applications, including skin moisturizers, osteoarthritis treatment, wound healing, adhesion prevention after surgeries, and artificial tissue engineering (Dicker et al. 2014; Fallacara et al. 2018; Lou et al. 2018).

**Figure 1.**
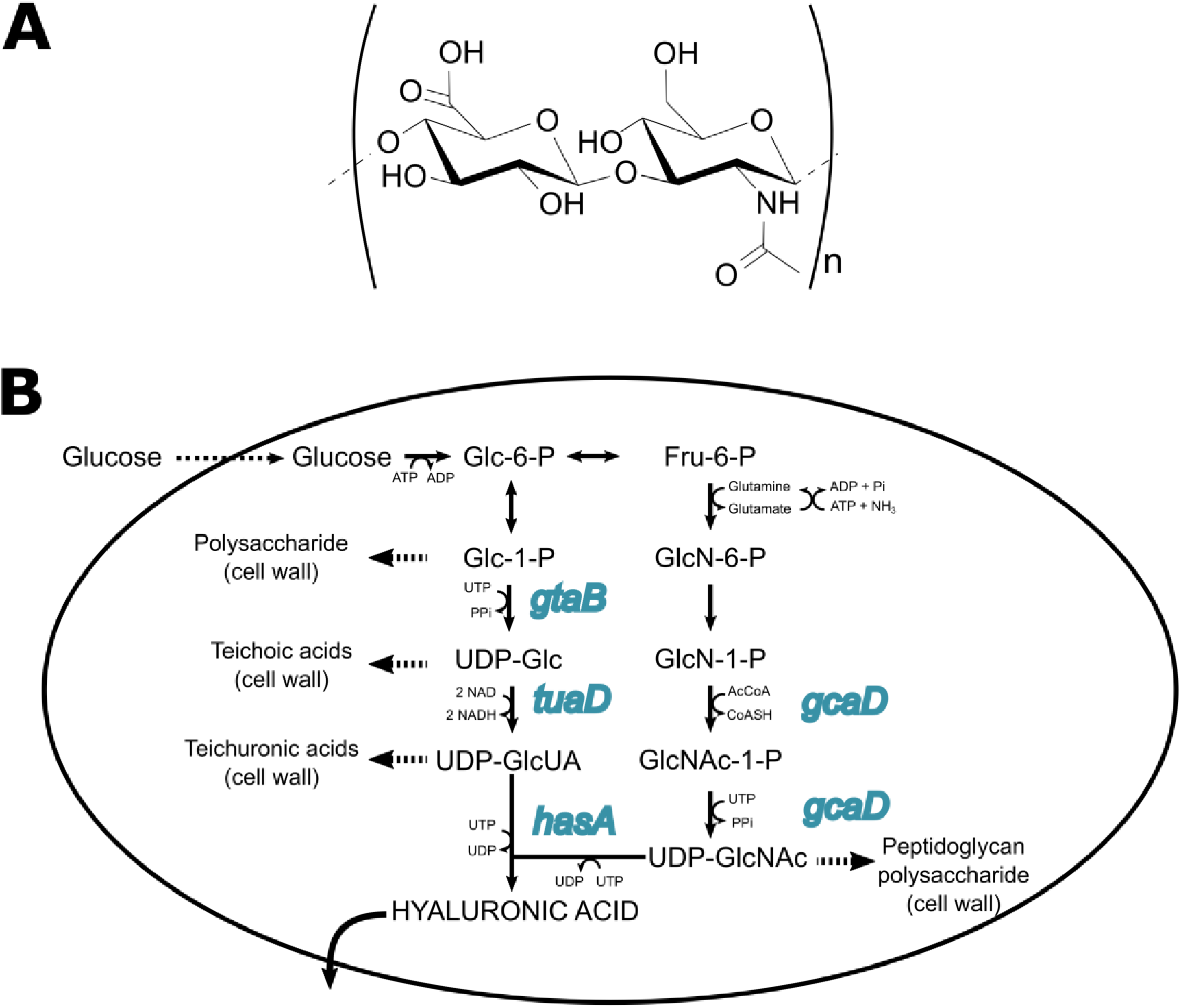
Hyaluronic acid structure. (A) representation of the repetitive disaccharide unit of hyaluronic acid, composed of β-1,3 and β-1,4-linked glucuronic acid and N-acetyl-D-glucosamine. (B) Biosynthetic pathway for the production of HA in engineering *B. subtilis* cells. Overexpressed genes are highlited.

HA was traditionally obtained from rooster combs, and today is mostly prepared by fermentation of *Streptococci* attenuated strains, such as Lancefield group A and C (DeAngelis et al. 1993; Kumari and Weigel 1997). In these microorganisms, HA is produced by a hyaluronan synthase (HasA), as a result of the alternative addition of GlcNAc and GlcUA units from their respective nucleotide sugars, UDP-GlcNAc and UDP-GlcUA, while the growing chain is exported outside the cell (Figure 1B).

The Streptococci-based HA raise safety concerns and are rejected by many consumers, since these microorganisms have the potential to produce exotoxins (Barnett et al. 2015; de Oliveira et al. 2016). A second problem is that Streptococci grow in complex culture media. These media are contamination-prone, generate waste with high organic loads, multiple operations are needed to isolate high quality HA, and may contain impurities not acceptable by regulatory agencies (Boeriu et al. 2013).

The HA global market for the cosmetic and pharmaceutical industries is estimated to be above 15 billion US$ in 2025 (Fallacara et al. 2018). Aside from these applications, HA and derivatives have extraordinary water binding capacities and are biodegradable, and are attractive candidates to be used in spill control, or in water absorbent pads, including diapers (Kurane and Nohata 1994). A large opportunity arises in the food industry, where the use of HA and derivatives is being explored as a component of functional foods and to build 3D frames along with collagen for cultivated animal-free meat, projected to be a trillion US$ market in 2040 (Ben-Arye and Levenberg 2019; de Souza et al. 2019; Oe et al. 2017; Zając et al. 2017).

To seize these opportunities, the cost effectiveness, safety and sustainability of the current manufacturing processes need to be improved.

In the last two decades, several attempts have been made to produce HA in safe microorganisms (Chien and Lee 2007b; Hoffmann and Altenbuchner 2014; Widner et al. 2005). *B. subtilis*, classified as a generally recognized as safe (GRAS) bacterium, is one of the most explored (Chien and Lee 2007a; Jia et al. 2013; Jin et al. 2016; Li et al. 2020; Westbrook et al. 2018; Widner et al. 2005; Zhang et al. 2016). Winder et al (2005) reported for the first time the heterologous production of HA in *B. subtilis*. They reached the upper limit of viscosity for stirred vessel fermentations, equivalent to ~7 g/L of HA, after 25 h using an engineered strain growing in a chemically defined salt-based medium (CDM). The medium composition was not disclosed, but a final cell density of 15 g DCW/L was informed. This is in agreement with reports indicating that most of the *B. subtilis* strains deficient in sporulation-specific sigma factors, can only reach DCWs between 5 and 15 g/L in CDM (Chen et al. 2013; Huang et al. 2003; Huang et al. 2004; Martınez et al. 1998; Pierce et al. 1992; Reuß et al. 2015; Wu et al. 2013; Yao et al. 2010). However, the production of HA in this host still faces many cost-related challenges. Only a few reports describe HA production in GRAS heterotrophic strains using CDM, but the VPs are in all the cases much lower than those reported for *Streptococci* -based commercial processes (de Oliveira et al. 2016).

The heterotrophic *B. subtilis* 3NA is a non-sporulating strain described in 2015 by Reuß and coworkers as a hybrid between *B. subtilis* 168 and its sub-species the W23 strain (Reuß et al. 2015). The strain shows a high growth rate in CDM, where it can reach cell densities of up to 75 g DCW/L in fed-batch cultures. This distinctive phenotype makes the 3NA strain an excellent candidate to be used as a host for the industrial production of biomolecules.

In modern fermentation, the use of CDM, containing only inorganic low-cost compounds is preferred (Stanbury et al. 2017). Based on the elemental composition and balance-mass equations, the components can be precisely dosed at quantities sufficient to obtain the product. In this way, costs reductions of raw materials in up to 80% can be obtained with a concomitant fall in the process waste (Dahod et al. 2010). In CDM, contamination events are reduced, fewer downstream operations are required, and the productivity is significantly more consistent, avoiding lot to lot deviations caused by regional or seasonal quality changes in the complex ingredients (Dahod et al. 2010; Reisman 2019; Stanbury et al. 2017).

The purpose of this study was to test a platform for the low cost and sustainable production of biomolecules. As a proof of concept, the robust *B. subtilis* 3NA strain was engineered to produce HA, a CDM formulated according to the elemental composition of *B. subtilis, and*. fermentation process with the maximum VP designed. A techno-economic analysis indicates that producing HA using this platform has remarkable benefits compared to the methods reported so far.

## Materials and methods

### Strains, plasmids and growth conditions

The *Escherichia coli* Top10 (Invitrogen) strain was used for general cloning purposes. *E. coli* strains were cultivated in Luria Bertani (LB) medium (10 tryptone, 10 g/L; yeast extract, 5 g/L; and NaCl 5 g/L) supplemented with 100 mg/L ampicillin or 50 mg/L kanamycin at 200 rpm and 37 °C.

*B. subtilis* 3NA cells were made competent by the method of Anagnostopoulos and Spizizen (Anagnostopoulos and Spizizen 1961). The *B. subtilis* 3NA strains carrying different constructions (Table S1), were used for HA production. For general purposes, these strains were grown in LB medium or Spizizen minimal medium at 200 rpm and 37 °C.

### DNA preparation, cloning and transformation

In order to obtain synthetic operons containing different combinations of genes of the HA biosynthetic pathways enzymes, all relevant genes, *i.e. has*A, *tua*D, *gta*B and *gca*B, were PCR amplified using the primers listed in the Electronic Supplementary MateriaL, Table S1, and genomic DNA from *S. zooepidermicus* ATCC6580, when amplifying *hasA*, or *B. subtilis* 3NA when amplifying *tua*D, *gta*B and *gca*B. Primers were designed in order to add a *Xba*I and/or *Nde*I restriction sites on the 5’ end and *Spe*I in the 3’ end of all genes in order to facilitate sequential cloning into the pTRG5 vector.

In order to provide strong ribosomal binding sites (RBS), the optimal consensus sequence of *B. subtilis* RBS 5’-AAGGAGG-3’ was included in the primer for amplifying *tua*D gene. The construction of the different operons is depicted in Figure S1. For the proof of principle, the IPTG-inducinle *P*_hyper-spank_ promoter was used to control the expression of the different operons. The constructed operons, in the pET24-derived plasmids, were digested with *Xba*I and *Spe*I and cloned into the *Nhe*I site of the pDR111 vector. The correct orientation was checked via PCR using a primer annealing in the inserted fragment a primer annealing in the *lac*I sequence present in the plasmid (Table S1). The resulting plasmid was integrated into the *amy*E locus of competent 3NA cells with selection for spectinomicyn resistance (50 mg/l^−1^).

#### High cell density fermentation process

Fed-batch fermentations were carried out in 3 L (New Brunswick Bio Flo 115, USA) fermenters. The temperature, stirring and pH were maintained at 37 °C, 1200 rpm and pH 7.0 (by addition of 30% NH_4_OH), respectively. The level of dissolved oxygen was controlled at 30 % of air saturation by changing the gas inflow (0.1 to 1 vvm) and the pure oxygen percentage when necessary. The seed culture was grown in batch mode with 40 g/L of glycerol to a final cell mass of ~17 g DCW/L. 10% vol/vol of the seed culture were diluted in a bioreactor containing the same medium supplemented with 1 g/L of glycerol and 0.1 mM IPTG. The feeding process was initiated when the glycerol present in the medium was exhausted.

Two different feeding strategies were used using a solution containing 800 gl^−1^ glycerol and 20 g/l MgSO_4_·7H_2_O. In the first one, the feeding rate (F, g/Lh) was determined by equation 1 (Lee 1996) in order to maintain the indicated specific growth rate (μ):

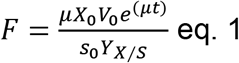

Where X_0_ is the initial biomass concentration (g/L), V_0_ is the initial volume (L), μ is the desired specific growth rate (h^−1^), S_0_ is the glycerol concentration in the feeding solution (g/L) and Y_X/S_ is the substrate yield (g/g). The second one consisted in an increase of glycerol from 2.5 to 9 g/L of the feeding solution in two hours, maintaining a constant feeding of 9 g/L·h until the end of the fed-batch fermentation process.

In both cases, production of HA was induced at the beginning of the fed-batch by adding IPTG at a final concentration of 0.1 mM. The supernatant was used to determine HA at the indicated points.

After the fermentation process, the cell culture was diluted in half in NaCl solution (final concentration of 2 M), and centrifuged at 5000 g for 10 min or the cells separated by microfiltration with a hollow fiber unit followed by ultrafiltration in a 750 KDa UF unit. For final formulation, HA was separated by ultrafiltration using a 0,1 μm MWKO hollow fiber cartridge followed by diafiltration to formulate the HA in pure water.

### Hyaluronic acid concentration and MW determination

HA samples were extracted from the fermentation broth at the times indicated. The cultures were centrifuged at 5000 *g* for 5 min. Amberlite IR120 (Sigma) was added to the supernatants until pH <4. The mixture was then centrifuged for 10 min at x 8000 *g* and NaOH was used to reach pH 7.0 of the resulting supernatants.

HA was precipitated with two volumes of ethanol and incubated at 4 °C for 1 h. The precipitate was collected by centrifugation at 5000 *g* for 10 min, and then redissolved in an equal volume of distilled water to measure the HA concentration via a modified Blumenkratz method (Blumenkrantz and Asboe-Hansen 1973). Briefly, a 200 μL sample was mixed with 1.2 mL of 2.5 g sodium borate in 1 L H_2_SO_4_ solution on ice. The mixture was boiled for 5 min and 12 μL methahydroxydiphenil 0.15% in 0.5% NaOH were added while vortexing the samples. Uronic content was determined by measuring absorbance at 520 nm. Commercial glucuronic acid 1 mg/mL was used as standard.

For molecular mass determination, further purification of the samples was performed by re-precipitating the samples with two volumes of ethanol. The samples were next resuspended, and dialyzed against water using a 100 KDa cut-off membrane (Sigma).

The weight-average molar mass and Mw distributions of HA were determined by high-performance size-exclusion chromatography (HPSEC) with on-line multi-angle light scattering (MALS), fitted with a K5 cell and a laser wavelength of 658 nm, and a refractive index detector. 100 μL of a 2 g/L sample were injected in the column [SB 806 M HQ columns (Shodex)] and eluted with 0.1 M NaNO_3_ containing 0.03% NaN_3_ at 0.5 mL/min. Solvent and samples were filtered through 0.1 μm and 0.2 μm filter units (Millipore), respectively.

### Techno-economic analysis

The simulation was made using the following assumptions:

The plant is located in the region of Rosario, Argentina. The total cost to start the business is 130 million dollars (US$) and includes the land, the facility with the building, all capital expenditures, working capital and all other expenses required to start the business (Electronic Supplementary Material, ESM_2.xlsx). The plant operates continuously for 330 days per year. Other infrastructure, like pilot plants, research laboratories, and sales offices space are not considered here. Financing was not included, and annual depreciation costs were equally distributed in 10 years. Annual maintenance rate of equipment was set as 2% of the direct fixed capital (DFC) cost.

Four production fermenters with a working capacity of 100,000 L each are used for HA production. A seed train is employed where the last seed fermenter transfers 10,000 L of an inoculum grown in batch mode to a final cell density of 15 g DCW/L. At the end of the fermentations, the cultures are transferred to an auxiliary tank, 5% of the broth is kept in the fermentor and fresh medium from reservoir tanks fed with a continuous sterilization system is added to start a new fermentation cycle. After three days of semi continuous work the fermentor is cleaned and steam sterilized. This turnaround takes 24 h, to calculate the duration of each fermentation cycle, the turnaround time is divided by the number of runs and proportionally assigned to the row “average fermentation cycle”. The turnaround time for the fermentor with complex media is assumed 10% longer because more time is required for cleaning than when using a CDM.

It is assumed that 96 % of fermentations are successful for CDM and 93% success is obtained for fermentations with complex media due to higher frequency of contaminations or other problems associated to complex media. HA titers are 6.8 Kg/m^3^ and the recovery yield of the downstream processing is 92%. The final product is HA with an average MW ~ 1 MDa.

Prices of the chemicals and consumables were obtained from local suppliers for quantities of at least one ton and salaries from a local survey. Technical grade glycerol was used, since it is locally available at large quantities from biodiesel producers and is 30% cheaper than glycerol pharma grade.

## Results

### *B. subtilis* 3NA was engineering for HA production

*Bacteria* naturally synthesize UDP-GlcUA and UDP-GlcNAc, a basic component of the peptidoglycan, also called murein, as intermediates for the assembly of cell wall structures (de Oliveira et al. 2016). Heterologous expression of *has*A in the *B. subtilis* 3NA strain is necessary and sufficient to confer to this host the capacity to produce HA. In addition, it has been shown that the over-expression of some of the native enzymes involved in the production of murein, and HA precursors, along with *has*A, increase the productivity (Figure 1B) (Jin et al. 2016; Widner et al. 2005).

To the best of our knowledge, HA has not been produced before in *B. subtilis* 3NA. Thus, derivative strains carrying *has*A from *S. zooepidemicus* (*has*A_Szo_) and increase copy number of the native *tuaD*, *gca*D and *gtaB* genes naturally present in the host were engineered. Three cell lines containing different IPTG-inducible expression cassettes inserted into the *amy*E locus were constructed. The first one (KCNHA8) expresses only *has*A_sz_; the second contains a bicistronic operon formed by *has*A_sz_ and an additional copy of the native *tuaD* gene (KCNHA9); and a third one (KCNHA0) with a four-gene operon (*has*A_sz_-*tuaD*-*gca*D-*gtaB* genes) (Electronic Supplementary Material ESM_1; Fig S1, Table S1).

The HA production of the three strains was tested in shake flask cultures using Spizizen minimal medium supplemented with 10 g/L of glycerol. Under the tested conditions, the co-expression of *has*A with the three auxiliary genes in the KCNHA10 strain gave the highest HA titer, producing more than twice the quantity obtained with KCNHA9 and four times the amount of the KCNHA8 cultures (Figure 2). No further productivity improvements were observed when the remaining genes of the pathways for the synthesis of UDP-GlcUA and UDP-GlcNAc (*pgi*, *gpm*, *glmM* and *glmS*) were over-expressed (data not shown).

**Figure 2.**
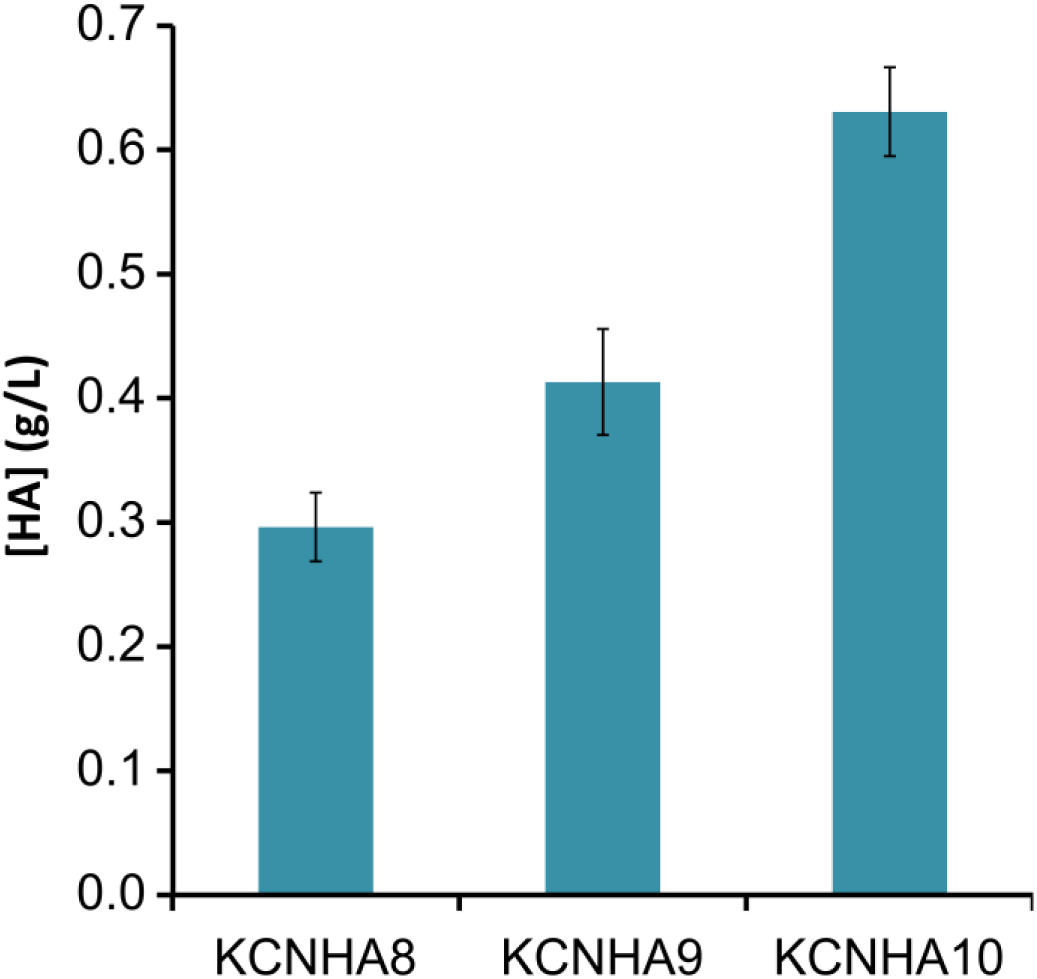
Titers of HA in batch cultures of *B. subtilis* 3NA derived strains grown in salt-based medium. KCNHA8, expresses hasA_Sz_ KCNHA9, expresses hasA_Sz_ and has an extra copy of *tuaD*, KCNHA10 expresses hasA_Sz_ and has extra copies of *tuaD*-*gca*D-*gtaB.*

#### Culture medium formulation

The efforts for lowering the manufacturing cost presented in this section were addressed to design a culture medium fulfilling the following constraints: (i) the medium should have the least possible amount of the cheapest raw materials, (ii) it should be based on inorganic salts to facilitate the purification of high quality HA and avoid effluents with a high load of organic matter, and (iii) the producing strain should grow at a high μ (specific growth rate, g/g·h) and the cells should produce HA at high q^P^ (specific productivity, mg/g of cells·h), to shorten the process time, in order to increase the VP (g of HA/·L h).

To build a given mass of cells, the quantities for each medium component were calculated based on the elemental composition of *B. subtilis* reported for carbon limited chemostat cultures (Dahod et al. 2010; Dauner et al. 2001).

The empirical formula for *B. subtilis* cell growing at 0.4 h^−1^ is C_1_N_0.221_P_0.020_S_0.006_ (Dauner et al. 2001), where carbon represents 48 % of the DCW, N is 12.3 %, P is 2.6% and S is 0.6 %. Because of its local availability at low prices, technical grade glycerol was used as a carbon and energy source. The yield coefficient (Y) for this carbon source, obtained from fed-batch fermentations in our laboratory, is 0.41 g/g, which means that, 2.44 g of glycerol (1g /0.41) are required to provide all the carbon contained to obtain one gram of dry *B. subtilis* cellular mass (DCW). Such amount of glycerol contains 0.94 g of C; from which 0.48 g (48 % x 1g) are converted, under aerobic conditions, in organic compounds that are part of the biomass, and the remaining C generates CO_2_ and energy (Dahod et al. 2010).

Because of their low cost, industrial grade H_3_PO_4_, and NH_4_OH were chosen as P and N sources, respectively. According to the mass ratio in the elemental formula, 0.026 g of P and 0.123 g of N, are needed for 1 g DCW of biomass; thus, 0.082 g of H_3_PO_4_ and 0.29 g of NH_4_OH were listed in the medium recipe.

Since NH_4_OH is toxic at high concentrations, 0.04 g/L was added in the form 0.1 mL of 30 % water solution. This quantity was calculated to accommodate the exact amount of potassium required to build 1 g DCW of biomass. K was provided by adding 0.023 g of KOH, the amount of base necessary to equilibrate the pH to 7.0. The remaining 0.99 mL of NH_4_OH were gradually fed during the course of the fermentation to maintain pH 7. In the same way, the quantities of sulfur and magnesium indicated in the elemental composition of *B. subtilis* biomass were provided as MgSO_4_ 7H_2_O. For the minor elements, low cost salts with high availability were used; also in amounts sufficient to provide the quantities reported in the biomass elemental composition analysis (Dahod et al. 2010). EDTA, a non-metabolizable chelating agent, was used instead of citrate; since this compound is used as a carbon source by the 3NA strain, causing the precipitation of divalent cations, which resulted in early growth interruption. The formula of this medium, CDM-1, designed to produce 1 g of DCW is shown in Table 1.

**Table 1.** Composition and cost of the CDM-1 medium, designed to obtain 1 g DCW/L of *B. subtilis*, supplemented with the additional quantities of Glycerol and NH4OH required or the biosynthesis of 7 g of HA

| Raw Materials | Price (US\$/kg) | g/L | US\$/L |
| --- | --- | --- | --- |
| Glycerol | 0.36 | 2.44 | 0.00088 |
| MgSO<sub>4</sub>·7H<sub>2</sub>O | 0.51 | 0.066 | 0.000034 |
| H<sub>3</sub>PO<sub>4</sub> | 0.89 | 0.082 | 0.000073 |
| NH<sub>4</sub>OH 28% (mL) | 0.41 | 1.03 | 0.00042 |
| KOH | 0.45 | 0.023 | 0.000011 |
| Na<sub>2</sub>-EDTA | 282 | 0.00033 | 0.000094 |
| Water (mL) | 0.00022 | 997 | 0.00022 |
| Na<sub>2</sub>[B<sub>4</sub>O<sub>5</sub>(OH)<sub>4</sub>]·8H<sub>2</sub>O | 0.45 | 0.0003 | 1.35E-07 |
| CoCl<sub>2</sub> | 9.52 | 0.0001 | 9.52E-07 |
| CuSO<sub>4</sub>·5H<sub>2</sub>O | 2.3 | 0.00021 | 4.83E-07 |
| FeCl<sub>3</sub>·6H<sub>2</sub>O | 7.63 | 0.0024 | 1.8312E-05 |
| MnCl<sub>2</sub> | 1.71 | 0.00073 | 1.254E-06 |
| Na<sub>2</sub>MoO<sub>4</sub>·2H<sub>2</sub>O | 12.31 | 0.00021 | 2.5851E-06 |
| ZnSO<sub>4</sub>·7H<sub>2</sub>O | 1.55 | 0.00033 | 5.0633E-07 |
| CaCl<sub>2</sub>·2H<sub>2</sub>O | 1.03 | 0.0025 | 2.575E-06 |
| Antifoam | 10.8 | 0.1 | 0.00108 |

On top of the components added to build the wished cell mass, extra amounts of C and N must be added for the synthesis of HA. For this, a yield coefficient of 0.31 g/g, determined by Don et al. (Don and Shoparwe 2010),) was used for glycerol, since the synthesis of HA requires significant amounts of energy. Thus, to obtain 7 g of HA, 22.1 g of glycerol (7/0.31) must be added to the medium. Based on the HA empirical formula, 0.24 g of extra N are needed to produce 7 g of HA, equivalent to 0.54 g of NH4(OH).

#### Fermentation process design

Having a CDM formulation defined, the goal of the fermentation process development described in this section was to achieve the highest possible VP. In this particular case, the titer for high MW HA (> 1 MDa) is limited by the viscosity of the broth to ~7 g/L, which is unfortunately quite low for a biomolecule intended to be used as a biomaterial. Thus, the only way to increase the VP was to reduce as much as possible the process time required to reach such a limiting titer.

The two time-dependent variables that govern the VP in this case. are (i) μ, which indicates how fast the cell factories can be created and (ii) q^P^, that indicates how efficient are those factories to produce HA.

Thus, a set of experiments was designed to determine how μ and q^P^ influence each other, to determine if the production of HA is growth associated or not. For this, the q^P^ value of cultures grown at different specific rates was determined. In all the cases, culture conditions feasible to be reproduced at large scale were used.

The fermentation time was set in less than 12 h, since that is the time employed to achieve the maximum VP reported for high molecular weight HA for *S. equi* (Kim et al. 1996). A fed-batch culture grown at a constant μ = 0.25 (~50% μ max) and with a starting cell density of 5 OD_600_, equivalent to 1.7 g DCW/L is expected to be near 30 g DCW/L after 11 h of exponential feeding. Under the mentioned constraints, and assuming an average q^P^ of 70 mg/g DCW h could be maintained throughout the fermentation, a process time of 11 h should provide more than 7 g/L of HA.

A seed culture of the KCNHA10 was grown in batch mode with 40 g/L of glycerol. When glycerol was consumed the final cell mass was ~17 g DCW/L. The seed culture was added to reach a final 10% V/V in CDM-30 (designed to produce 30 g of DCW) with 1 g/L of glycerol and 0.1 mM IPTG to twin bioreactors. After 15 min, glycerol was exponentially fed to keep μ at 0.25 h^−1^ in the first reactor and at 0.35 h^−1^ in the second one.

Figure 3A and B show that in both experiments the final cell density was near 30 g DCW /L, indicating that the medium formulation was sufficient to build the expected mass of cells, but none of the experiments reached the expected HA titer. The culture grown at 0.25 h^−1^ produced ~3 g/L of HA and q^P^ was 23.03 ± 1.04 mg of HA/gDCW h (Figure 4A). The culture grown at 0.35 h^−1^ was stopped when keeping the dissolved oxygen above 20% was no longer possible and excessive foam was formed, indicating a progressive cell lysis. At that point 3 g/L of HA were accumulated and q^P^ was 52.67 mg ± 2.54 mg of HA/gDCW.h (Figure 4B). We observed in batch experiments q^P^ values above 80 mg of HA/gDCW.h when the cells were grown in excess of glycerol at μ max. Thus, a positive correlation between q^P^ and μ exists, at least in the range between 0.25h^−1^ and μ max.

**Figure 3.**
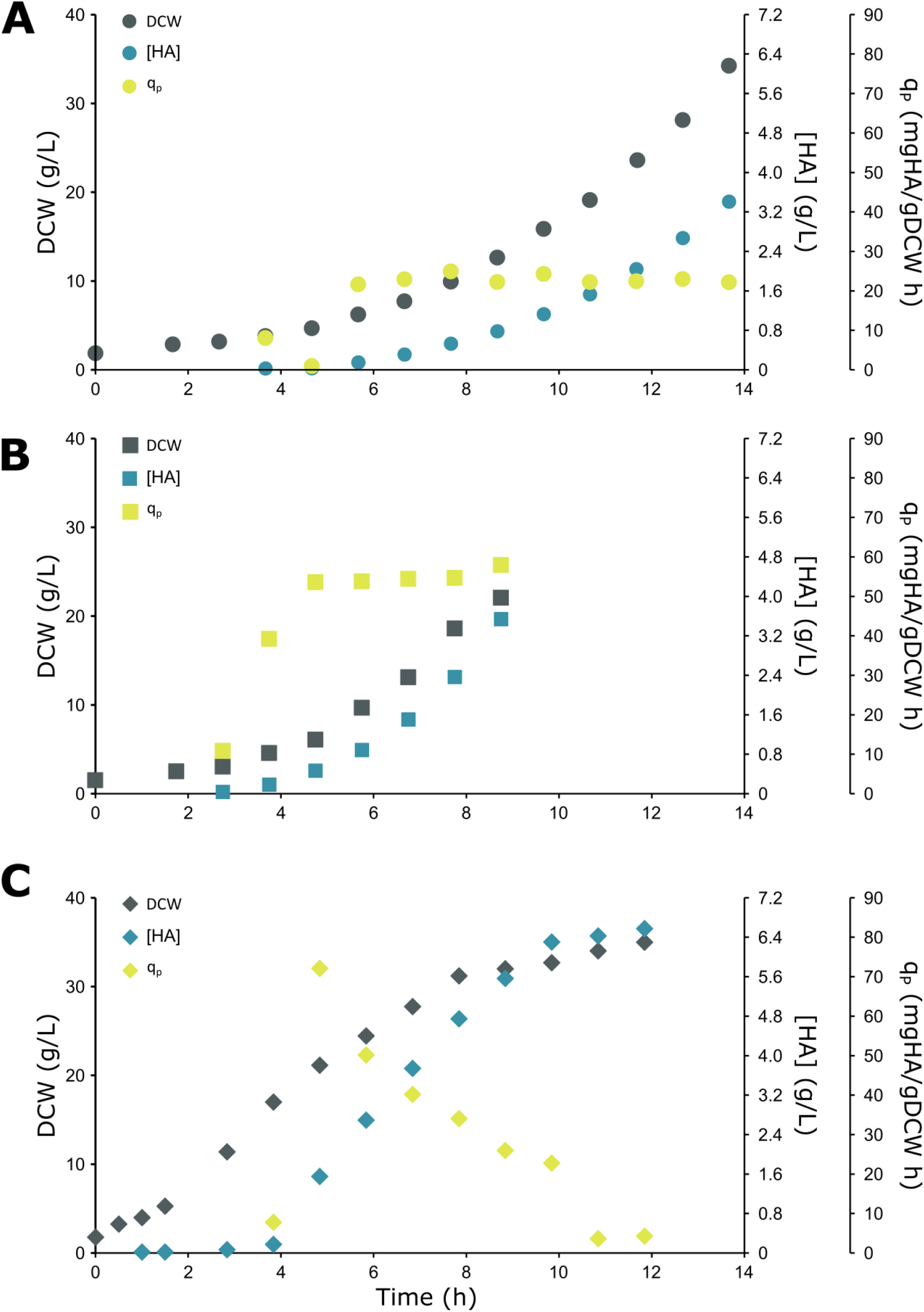
Biomass and HA production of KCNHA10 strain grown in CDM-30. (A) exponential feeding to keep μ = 0.25 h^−1^. (B) exponential feeding to keep μ = 0.35 h^−1^·(C) constant feeding rate of 9 g/L·h.

**Figure 4.**
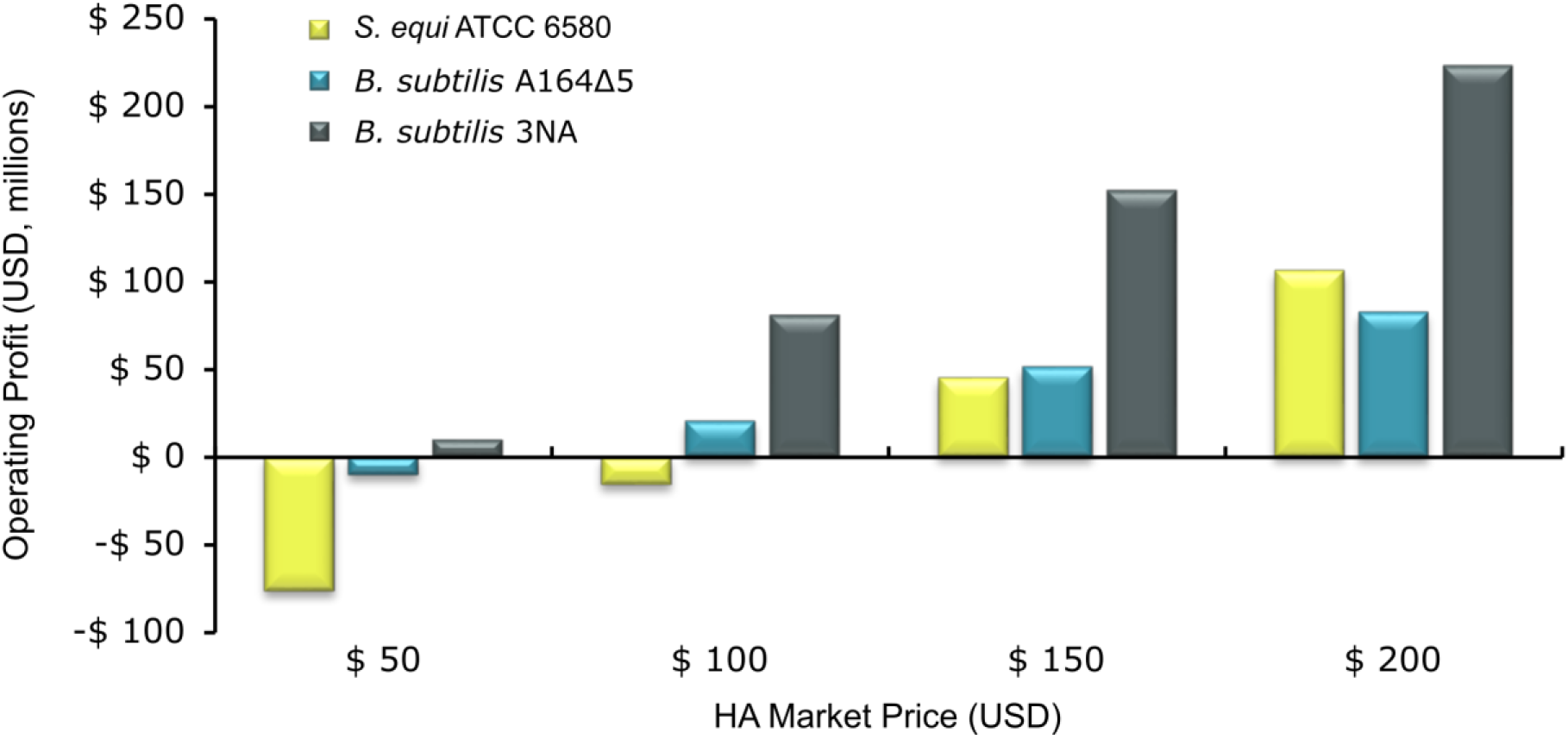
Sensitivity analysis of the operating profit of each process exposed to the HA market price.

Based on the result of the analysis, the preferred fermentation strategy was to maximize μ, since this variable in a CDM with limited carbon can be precisely controlled by increasing the feeding rate of the carbon source. A feeding of 9 mL/L·h was chosen, since this is the maximum value that can be later reproduced in large scale fermentation, and one hour later cells were growing at 0.53 h^−1^, the value for μ_max_ observed in batch cultures with excess of glycerol. As expected, q^P^ reached ~80 mg/g·h and the two variables converged to give the highest possible value VP for the system. The cells grew at 0.53 h^−1^ until the point where the glycerol uptake by the biomass present in one liter of the culture was large enough to consume all the glycerol and the feeding rate matched the uptake by the cell mass. From that point and to the end of the fermentation μ and q^P^ started to decrease and the biomass showed the expected linear growth.

Figure 3C shows that HA average titers near 7 g/L were consistently obtained after feeding glycerol for 10.5-11 h and that the final average cell density of 29.4 g DCW/L was achieved.

The HA obtained in CDM-30 medium was easily purified by (i) separating the cells and large cell debris with microfiltration; and (ii) separating trace of protein and DNA fragments from lysed cells and other low MW compounds using ultra and diafiltration steps. HPSEC-MALS analysis of purified HA obtained from fed-batch fermentation process performed at constant feeding rate of 9 g/L·h of glycerol showed a main peak of 9.53 10^5^ Da (± 3.53%) with a dispersion of 1.19 (± 4.54%).

#### Cost analysis

The techno-economic analysis presented below illustrate the potential impact of the process described in this work on the HA market. As a reference, the expected costs of two other methods extracted from the literature are included, when data is not provided the values were assumed to be the same as the process described here:

Table 2 Models the operating profit for the large scale HA manufacturing in a plant with a working capacity of 400 m^3^, and compares the process described here with two others. The first one, using *S. equi* grown in a complex medium, was chosen as an example of a typical Streptococci-based process. In this case, the final HA titer and fermentation time are similar to those of the method described here, which is reflected in the VPs. However, the cost of the culture media is more than five times higher than the CDM designed here, mostly due to the cost of tryptone required by the auxotrophic strain, and the HA purification.

**Table 2.**
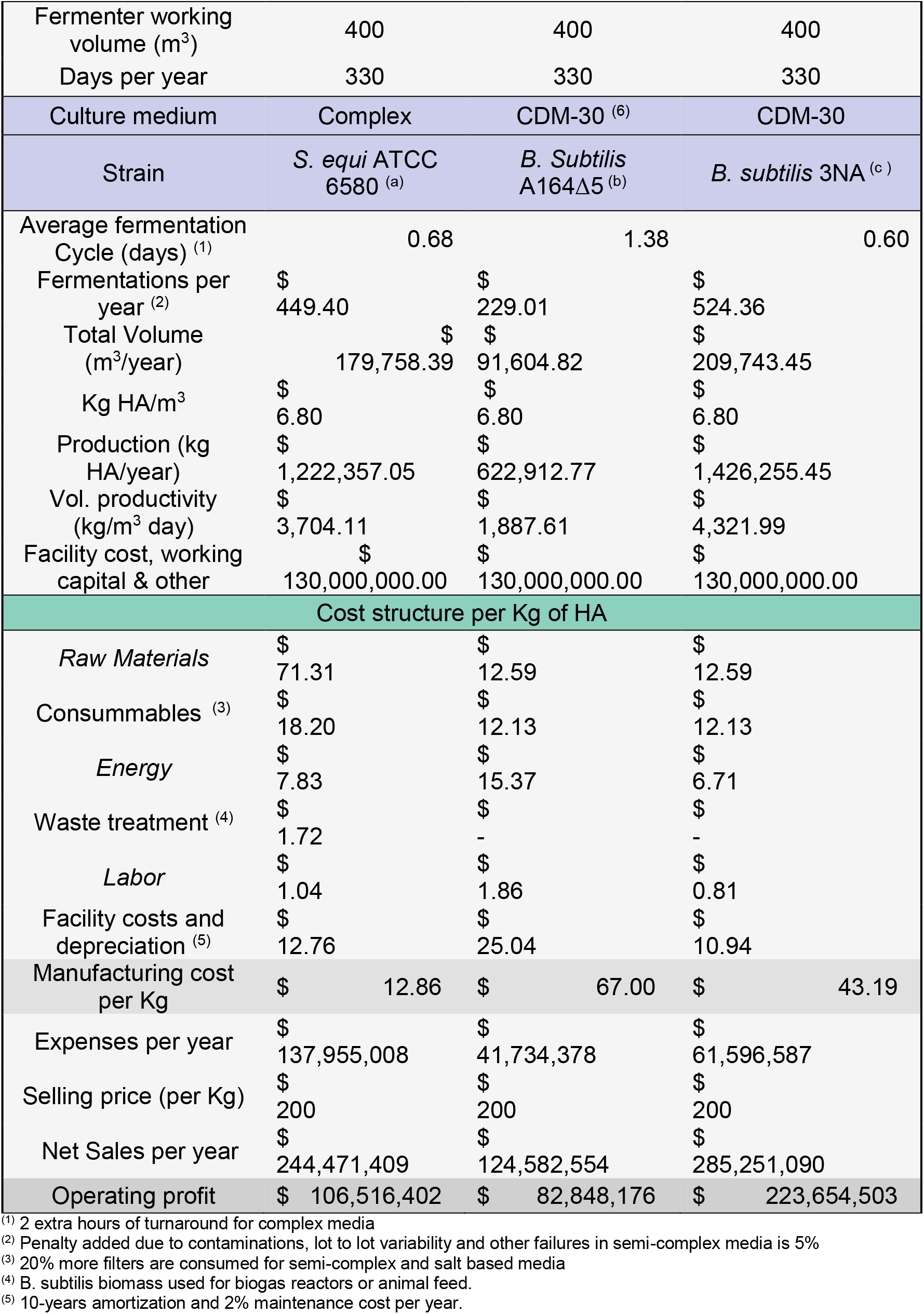
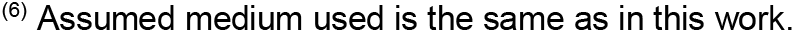
Techno-economic analysis for simulated HA manufacturing at 400 m^3^ scale using the process presented in this work, and two published processes. In the last two cases, some data was not available and the Table was completed with data from other sources, including our laboratory.

|  |  |  |  |
| --- | --- | --- | --- |
| Fermenter working volume (m<sup>3</sup>) | 400 | 400 | 400 |
| Days per year | 330 | 330 | 330 |
| Culture medium | Complex | CDM-30 <sup>(6)</sup> | CDM-30 |
| Strain | *S. equi* ATCC 6580 <sup>(a)</sup> | *B. Subtilis* A164Δ5 <sup>(b)</sup> | *B. subtilis* 3NA <sup>(c)</sup> |
| Average fermentation Cycle (days) <sup>(1)</sup> | 0.68 | 1.38 | 0.60 |
| Fermentations per year <sup>(2)</sup> | \$ 449.40 | \$ 229.01 | \$ 524.36 |
| Total Volume (m<sup>3</sup>/year) | \$ 179,758.39 | \$ 91,604.82 | \$ 209,743.45 |
| Kg HA/m<sup>3</sup> | \$ 6.80 | \$ 6.80 | \$ 6.80 |
| Production (kg HA/year) | \$ 1,222,357.05 | \$ 622,912.77 | \$ 1,426,255.45 |
| Vol. productivity (kg/m<sup>3</sup> day) | \$ 3,704.11 | \$ 1,887.61 | \$ 4,321.99 |
| Facility cost, working capital & other | \$ 130,000,000.00 | \$ 130,000,000.00 | \$ 130,000,000.00 |
| Cost structure per Kg of HA |  |  |  |
| *Raw Materials* | \$ 71.31 | \$ 12.59 | \$ 12.59 |
| *Consummables* <sup>(3)</sup> | \$ 18.20 | \$ 12.13 | \$ 12.13 |
| *Energy* | \$ 7.83 | \$ 15.37 | \$ 6.71 |
| *Waste treatment* <sup>(4)</sup> | \$ 1.72 | \$ - | \$ - |
| *Labor* | \$ 1.04 | \$ 1.86 | \$ 0.81 |
| *Facility costs and depreciation* <sup>(5)</sup> | \$ 12.76 | \$ 25.04 | \$ 10.94 |
| Manufacturing cost per Kg | \$ 12.86 | \$ 67.00 | \$ 43.19 |
| Expenses per year | \$ 137,955,008 | \$ 41,734,378 | \$ 61,596,587 |
| Selling price (per Kg) | \$ 200 | \$ 200 | \$ 200 |
| Net Sales per year | \$ 244,471,409 | \$ 124,582,554 | \$ 285,251,090 |
| Operating profit | \$ 106,516,402 | \$ 82,848,176 | \$ 223,654,503 |

<sup>(1)</sup> 2 extra hours of turnaround for complex media
<sup>(2)</sup> Penalty added due to contaminations, lot to lot variability and other failures in semi-complex media is 5%
<sup>(3)</sup> 20% more filters are consumed for semi-complex and salt based media
<sup>(4)</sup> *B. subtilis* biomass used for biogas reactors or animal feed.
<sup>(5)</sup> 10-years amortization and 2% maintenance cost per year.
<sup>(6)</sup> Assumed medium used is the same as in this work.

For the second comparison, part of the data was obtained from the work of Widner et al (2005), reporting for the production of HA in a prototrophic *B. subtilis* strain grown in a salt based medium. The culture was incubated for 25 h, but the medium composition is not described and they report a HA final titer “in the multigram range”. Even assuming that the process is terminated when titers reached ~7 g/L, the technical limit for high MW HA production, and that the cost of the medium used is as low as the one designed for this work, the expected VP would be 56% lower than the one projected for the method described here (Table 2). The difference is caused by the extended process time, which directly reduces the VP and therefore increases the cost (Ferreira et al. 2018). The sensitivity analysis of Figure 4 is based on the techno-economic analysis of Table 2. At the current HA bulk market price of 200 US$/Kg, the process presented here is 115% more profitable than the *Streptococci*-based processes used today. If the price drops to 150 US$, the operating profit would be 247% higher, while at prices below 114 US$ the *S. equi* process would not even reach the breakeven point. In the former scenario, producing HA with the 3NA strain would triplicate the operating profit obtained with the *B. subtilis* 165 strain in 25 h (Widner et al. 2005).

## Discussion

The goal of this work was to design from the ground a bioprocess to manufacture high molecular weight HA in a GRAS microorganism, analyzing every step where savings can contribute to achieve the lowest possible cost. For high MW HA, titers above 7 g/L cannot be obtained due to the constraint imposed by the viscosity of the broth and the consequent rapid drop of the oxygen transfer capacity (Boeriu et al. 2013; Fallacara et al. 2018). Thus, efforts were focused on

i. Engineering a prototrophic, fast growing and GRAS HA producing strain.
ii. Designing a CDM with the least possible cost.
iii. Finding process conditions where the producing strain cultured in the designed medium reaches the maximum VP.

The *B. subtilis* 3NA strain was reported to grow to high cell densities in CDM (Reuß et al. 2015). In our laboratory, a notable capacity to grow fast in CDM was observed. These features make the 3NA an ideal host to be explored for the development of high VP fermentations using inorganic salts as raw materials, the type of bioprocess needed for manufacturing bio-commodities like HA.

Strains harboring the heterologous *has*A_Sz_ gene produced more HA when additional copies of three native genes involved in the production of HA precursors were overexpressed. Similar results were shown in other *Bacillus* stains (Jin et al. 2016; Widner et al. 2005), suggesting that the carbon flow through biosynthetic pathways of GlcNAC and GlcUA is at least similar. This is not surprising considering that the gene clusters are highly conserved.

For microorganisms with a long history of industrial use, the culture media were empirically shaped throughout decades, but contain a large excess of the elements required to build the required mass of cells and products. CDMs facilitate downstream processing with the consequent cost reduction, minimize contamination events, and improve the reproducibility of the fermentations (Dahod et al. 2010; Stanbury et al. 2017).

Here, for a rational design of a CDM, the amount of carbon source to be added was first calculated based on the yield coefficient. Oxygen is provided by the air flow and limited by the oxygen transfer rate (OTR) of the bioreactor. For all the other elements, the quantity of each inorganic salt to be added was calculated according to the elemental formula of a *B. subtilis* to build one gram DCW of cells (Table 1). To reach the least possible cost, the cheapest inorganic salts and carbon source were chosen. The design of the CDM medium is flexible and allows the users to customize the formulation by keeping the relative ratio of the components. According to the final cell density of *B. subtilis* 3NA required for a given bioprocess, the amounts of each component can be determined using the Electronic Supplementarry Material, ESM_2,0020adding in each case the amounts of C and other elements needed to support de production of the target bioproducts.

Rigorous comparisons of bioprocess performances supported by costs analysis are scarce in the literature. Often, a fermentation process is considered successful based on an isolated piece of information, like the product titer or the use of some waste as a low cost material, ignoring industrial needs, like operating parameters and even the process time. The VP, where the process time is taken into account, along with the cost of all the raw materials employed, are the metrics that define the success of a fermentation process.

For the rational design of a fermentation process, the interactions between μ and q^P^, the variables that define the VP for a given volume were studied. The two variables may have a positive correlation, like in the case of proteins expressed using the GAP promoter in *P. pastoris*, or β-galactosidase from *E. coli* in *B. subtilis*; where the expression of the target proteins is growth associated. In other cases, the product can be non-grow associated like certain secondary metabolites in the genus *Streptomyces,* mixed growth associated, or the correlation can be negative as well (Looser et al. 2017; Martínez et al. 1998; Sakthiselvan et al. 2019).

In *B. subtilis 3NA*, a positive correlation between μ and q^P^ was found, in the range of μ between 0.25 and 0.53, similar results were found in *S. zooepidermicus* grown in complex media in batch experiments (Don and Shoparwe 2010). The culture conditions that maximized the values of μ and q^P^, were determined, limited by the constraints that guarantee a smooth transfer to large scale bioreactors. To the best of our knowledge the VP achieved is the highest reported for HA with MW ~1 MDa using a GRAS host.

In summary, the CDM medium, the capacity of the *B. subtilis* 3NA strain to reach high cell densities and the designed the fermentation process, provide the basis to obtain a remarkable cost reduction of high MW HA manufacturing at large scale.

- First, the prototrophic phenotype of the *B. subtilis* 3NA allows for the use of CDM with a cost estimated to be 80% lower than the media used for HA production in Streptococci.
- Second, reaching high cell density in CDM provides the 3NA strain an advantage over any other *B. subtilis* strain described so far, since more producing cells shorten the process time to reach the limiting titer of HA and therefore increasing the VP. Widener and co-workers (2005) produced in a derived *B. subtilis* 165 strain 7 g/L of HA in 25 h, while the data presented here showed that the 3NA strain produced the same amount in 11 h; resulting in a VP more than 100% higher (Table 2 and Electronic Supplementary Material ESM_2)

HA is a biodegradable polymer with remarkable properties to be used in many industrial applications, but the current price of this biomaterial is a barrier for its widespread adoption. As the patents protecting production methods expire, the price of bioproducts naturally tend to decrease due to the arrival of new competitors. The advantages expected from the platform described in this work are exposed in Table 2 and in the sensitivity model of Figure 4. The platform presented here dramatically reduce the current manufacturing prices in 80% making available industrial quantities of HA as a food ingredient among other uses.

The *B. subitlis* 3NA strain, the CDM medium and the fermentation process compose a freely available platform that could be used in principle for the safe and environmentally friendly production of other biomaterials in a cost-effective manner. The guidelines of supplementary Material can be used as a starting point and the materials are available upon request.

## Conclusions

An environmentally friendly platform based on: (i) *B. subtilis* 3NA as a new chassis strain, (ii) a cost-effective chemically defined culture medium, and (iii) a fermentation process with a remarkable VP; was developed and validated by producing high quality HA at laboratory scale.

A techno-economic analysis indicates that a large scale manufacturing facility would be more than 100 % more profitable than with the methods reported so far for the production of HA with MW ~1 MDa. Scaling up the process described in this work may facilitate the use of HA as a food ingredient or as biomaterial for other purposes.

The IP free platform presented here can be applied to develop cost-effective and safe manufacturing methods for other biomolecules.

## Ethical statement

This article does not contain any studies with human participants or animals performed by any of the authors.

## Acknowledgment

The authors want to acknowledge Juan Diego All, Mauro Nicolás Cerchi, and Claudio Exequiel Cansino Quispe for their technical assistance. This work was partly supported by Glyco@Alps (ANR-15-IDEX-02).

